# Improved BOLD Detection with Sliced Inverse Regression

**DOI:** 10.1101/2024.02.21.581434

**Authors:** Andrew Lizarraga, Katherine Li

## Abstract

Functional magnetic resonance imaging (fMRI) has been effective in linking task-related brain responses to changes in blood oxygen level density (BOLD). However, its reliance on BOLD measurements makes it vulnerable to artifacts and false-positive signals. Commonly, researchers use many regressors in a General Linear Model to filter true signals, but this adds noise and complicates interpretation. In this paper we suggest using Sliced Inverse Regression (SIR) to simplify covariate dimensionality and identify relevant regressors. We compare a general linear model applied to both original and SIR-adjusted data, demonstrating that SIR improves signal detection, reduces noise, and yields statistically significant results even with conservative measures.

## 1 Introduction

The use of Blood Oxygen Level Density (BOLD) Jbabdi (2014) is widely studied and frequently used measure of activity in the brain. Typically the experimenter will have a subject in the scanner and measure brain activity at rest and measure BOLD response when the subject is given an activity such as identify an object on the screen or listen to some audio. When comparing the oxygen level saturation in the brain during certain responses, the MRI scanner can pick up these signals and associate them to certain regions (voxels) in the MR image. To determine signals that are statistically significant, researchers commonly use a General Linear Model (GLM) Monti (2011). However, BOLD responses are especially sensitive to artifacts such as overall blood pressure changes, motion artifacts due to head movement in the scanner, fluid movement (in particular: Cortico Spinal Fluid, CFS), ect. Jbabdi (2014) Monti (2011). This has led to a variety of methods including noise modelling in addition to the MRI scan. It’s suggested in Lund et al. (2006) that modeling cardiac and respiratory rate is needed in order to reduce these artifacts in BOLD detection. Other researchers include noise modeling along with specific experimental designs in order to mitigate noise artifacts Razavi et al. (2003-12). However, these methods can only be used a priori and don’t support efforts on early open-access data that aggregated fMRI scans prior to proposals in Lund et al. (2006) Razavi et al. (2003-12) Monti (2011). The efforts of this paper are to provide tools to assist GLM analysis of BOLD under non-ideal conditions.

The GLM in practice typically involves data points *X* ∈ ℝ^*p*^ and response signals *Y* ∈ ℝ, for each voxel in (*i, j, k*) ∈ ℝ^3^. We view *X* as consisting of parameters for drift signals during resting state and active state. The idea is that the experimenter doesn’t know when they will pick up a BOLD signal, so multiple drift terms are used. However it is often the case that not all regressors are indictive of BOLD and instead are correlated with artifacts that are not of use in BOLD studies Gorno-Tempini et al. (2002) Razavi et al. (2003-12) Lund et al. (2006). To counter this, we apply techniques from the Sliced Inverse Regression (SIR) Li (1991) Coudret et al. (2014) literature in order to reduce the dimensionality of *X* given the signal response we recorded in *Y*. This has the advantage that we select the regressors of *X* that correspond to statistically significant effects in *Y*. Before going into analysis, we brief the reader on SIR theory and literature that is relevant to this study.

### 1.1 Background

In this section, we brief the reader on sliced inverse regression. We start with the data *X* and its corresponding measured response *Y*. The goal of regression usually is to determine 𝔼 [*Y* |*X* = *x*]. Countless techniques exist for this task, including OLS regression, PCR, norm-regularized regression, and PLSR, to name a few. But these methods are limited in that they assume the relationship between *X* and *Y* is linear. Therefore, when relationships between variables are more complex, such as with fMRI data, this assumption breaks down, creating a need for new methods that can better treat nonlinear data dependencies.

One method that addresses this problem is sliced inverse regression. Although nonlinearities make the analyses more complicated for both forward and inverse regression, it is less of a concern for inverse regression. Furthermore, sliced inverse regression can act as a dimension reduction technique, simplifying *X* into a lower dimensional representation spanned by the effective dimension reducing directions (e.d.r’s) Li (1991). These advantages make sliced inverse regression an effective way to treat both higher dimensional and nonlinear data, especially when combined with other data reduction techniques.

Sliced inverse regression works as follows. Given the model 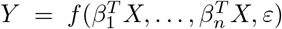, the conditional expectation 𝔼 [*X*|*Y* = *y*] is first determined. Then, from performing a general eigenvalue decomposition of the covariance matrix of this conditional expectation, 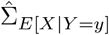, with respect to the covariance matrix of *X*, 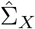, the eigenvectors *β*_*j*_ are considered as the e.d.r. directions Li (1991). While there is no linearity assumption on the relationship between *X* and *Y, X* is assumed to satisfy the linear design conditions, meaning that the regression of *X* on 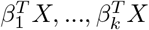, must be linear. Although this presents some limitations to this technique, it in fact relaxes the assumptions of normality in *X* that pervade many popular regression methods.

A primary aim for BOLD analysis is to remove regressors that don’t significantly affect the response variable. Because SIR takes into account the response when dimension-reducing *X*, we consider it to be a good candidate for our analysis. We focus on the active state task and leverage SIR to determine the regions of activation in the brain. In particular we focus on the auditory cortex Purves et al. (2001).

### 1.2 Related Work

There is a sparse collection of literature of SIR methods applied to fMRI, but notably, Tu et al. (2018) proposed Sliced Inverse Regression Decoding Analysis (SIR-DA) as a tool for fMRI. SIR-DA utilizes sliced inverse regression (SIR) for estimating effective dimension reduction (e.d.r.) directions, especially when the relationship between fMRI data and class labels is not clearly defined. It incorporates Singular Value Decomposition (SVD) with SIR to handle high dimensional data, specifically when the number of features, *p > n*, the sample size, and it may be more difficult to perform an eigenvalue decomposition due to the singularity of the covariance matrix. The SVD-SIR method then becomes integrated into the pattern classifier and it can effectively detect the signal even when additional noise is added to the response.

Tu et al. (2015) demonstrated on simulated data that SIR is more effective in identifying brain activation and predicting outcomes compared to support vector regression (SVR) and partial least square regression (PLSR), irrespective of the number of slices used in the inverse regression in general and more so when there is a strong signal to noise ratio in the fMRI data. Also, when applied to real fMRI data from a pain stimulation experiment, PC-SIR was able to identify pain-related brain regions and predict pain perception more accurately than SVR and PLSR. The outperformance of SIR indicate that it is a viable method for handling fMRI data that may warrant further exploration. In this paper, we apply a combined SIR and GLM approach, given that this is the most standard fMRI analysis. We are aware that there may be slower adoption of this method, since there isn’t thorough mention of it in the previous literature. Additionally, previous studies were not conducted on the same open-access data, making it difficult for authors to validate the findings. However, we show that applying SIR with GLM is sufficient for detecting relevant BOLD signals and actually can outperform other methods even on easily available open-access datasets. We hope to ease verification of our results by doing so.

### 1.3 Contributions

In this paper we make the following contributions:

- We provide an open-access analysis of SIR methods applied to fMRI
- We demonstrate that SIR methods are suitable for active state fMRI and in particular reveal auditory-associated regions of the brain that are anatomically valid and inline with current literature.
- We show that SIR dimension reduction on BOLD regressors leads to better signal detection and reduction of noise/artifacts then applying GLM methods alone.

## 2. Methodology

Note that, in BOLD, signal response is a times series in each voxel comprising the MR image for the brain. This make the response multi-variate and consequently we cannot directly apply SIR methods. However we can index on each voxel and consider the response to be a univariate signal and apply SIR iteratively across each voxel. This is sensible for BOLD since the MR image is constructed in a per voxel basis with measurements taken independently of each other Monti (2011).

Lastly, we determined the number of e.d.r. directions based on the amount of variability in our data that the directions could explain. Specifically, we chose a cutoff percentage of 95%, corresponding to 12 e.d.r directions, since there is drop-off in the amount of variability explained beyond this number of directions. We slice the data into 96 parts for computational ease. Since SIR is generally robust to the number of slices used Tu et al. (2018), we believe this is a reasonable approach.

The algorithm for the process is described as follows:

The Multi-SIR algorithm represents an adaptation of the existing SIR algorithm, tailored specifically for scenarios where the response is composed of independent signals. This is particularly relevant in our case, considering the nature of fMRI sampling Monti (2011). We operate under the assumption that certain artifacts, such as head motion, blood pressure variations, and breathing patterns, do not contribute to the BOLD signal, but rather constitute noise picked up by the regressors.

Our assumptions around noise have been corroborated in Sec. 3, where our analysis demonstrates that the GLM applied to data corrected by the SIR algorithm shows a high resilience to noise. Despite the presence of noise, the analysis successfully detects active signals, even under the application of stringent statistical tests. This outcome reinforces the effectiveness of the Multi-SIR approach in isolating true neural activity from noise in fMRI data.

### Algorithm 1

Multi-Index Sliced Inverse Regression (Multi-SIR)

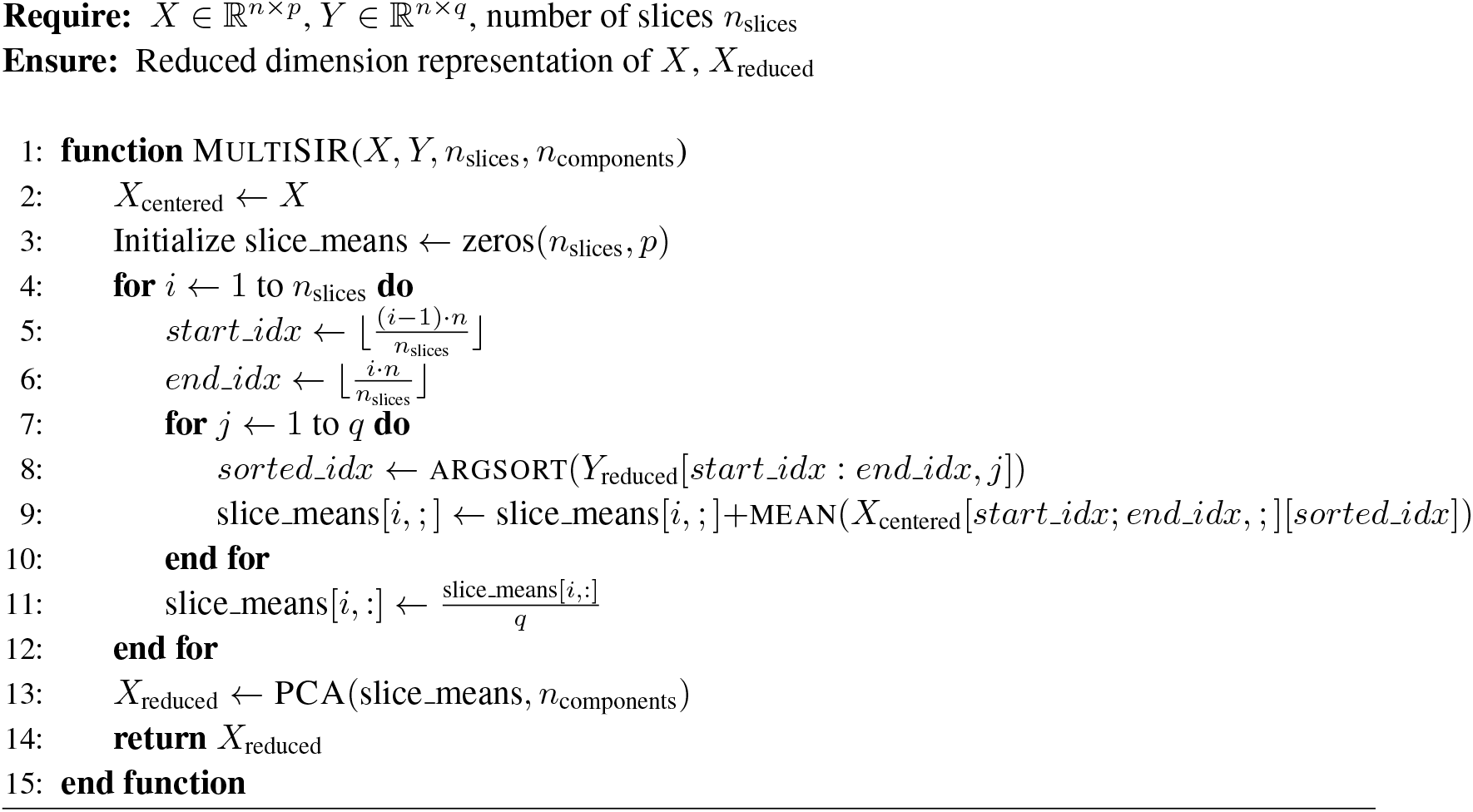

## 3. Experiments/Data Exploration

### 3.1 Data

As outlined in Sec. 1.1, the primary objective of a GLM is to identify voxels that exhibit a statistically significant response during active and resting states of the subject. This information is crucial for pinpointing brain regions that become active during specific tasks, thereby enabling researchers to correlate particular brain areas with specific activities. In our study, our focus is to locate the brain regions activated by auditory stimuli. Ideally, we expect to see aggregation of BOLD activation signals in these highlighted areas. We show that our Multi-SIR methodology with GLM is more inline with highlighting activity auditory cortex than the standard GLM.

We utilized an openly available dataset from the Nilearn Python library *Nilearn* python library Abraham et al. (2014). This dataset comprises 96 brain scans of a single subject from one session. Each scan has a repetition time (TR) of 7 seconds. The session included alternating active and rest periods, each lasting 42 seconds. During active periods, auditory stimulation involved the presentation of bi-syllabic words through headphones at a rate of 60 beats per minute. We specifically used data from the image file: fM0023 004. Detailed information on data processing and extraction is available through this GitHub link: https://github.com/drewrl3v/fmri-sir.

### 3.2 Experimental Results

We run a standard GLM against a GLM with SIR adjusted input data to learn a first order model. Detailed implementations are made publicly available, via the following GitHub repo: https://github.com/drewrl3v/fmri-sir. We then perform standard statistics seen in the fMRI community in order to gauge the behavior of these two models Monti (2011). As we see in Fig. **1** (4.3), the left column depicts the standard GLM with a variety of statistical tests indicating significance, while the right column is the detected response of the SIR adjusted GLM. Each row of Fig. **1** (4.3) consists of the following statistical tests used throughout fMRI literature:

**Figure 1.**
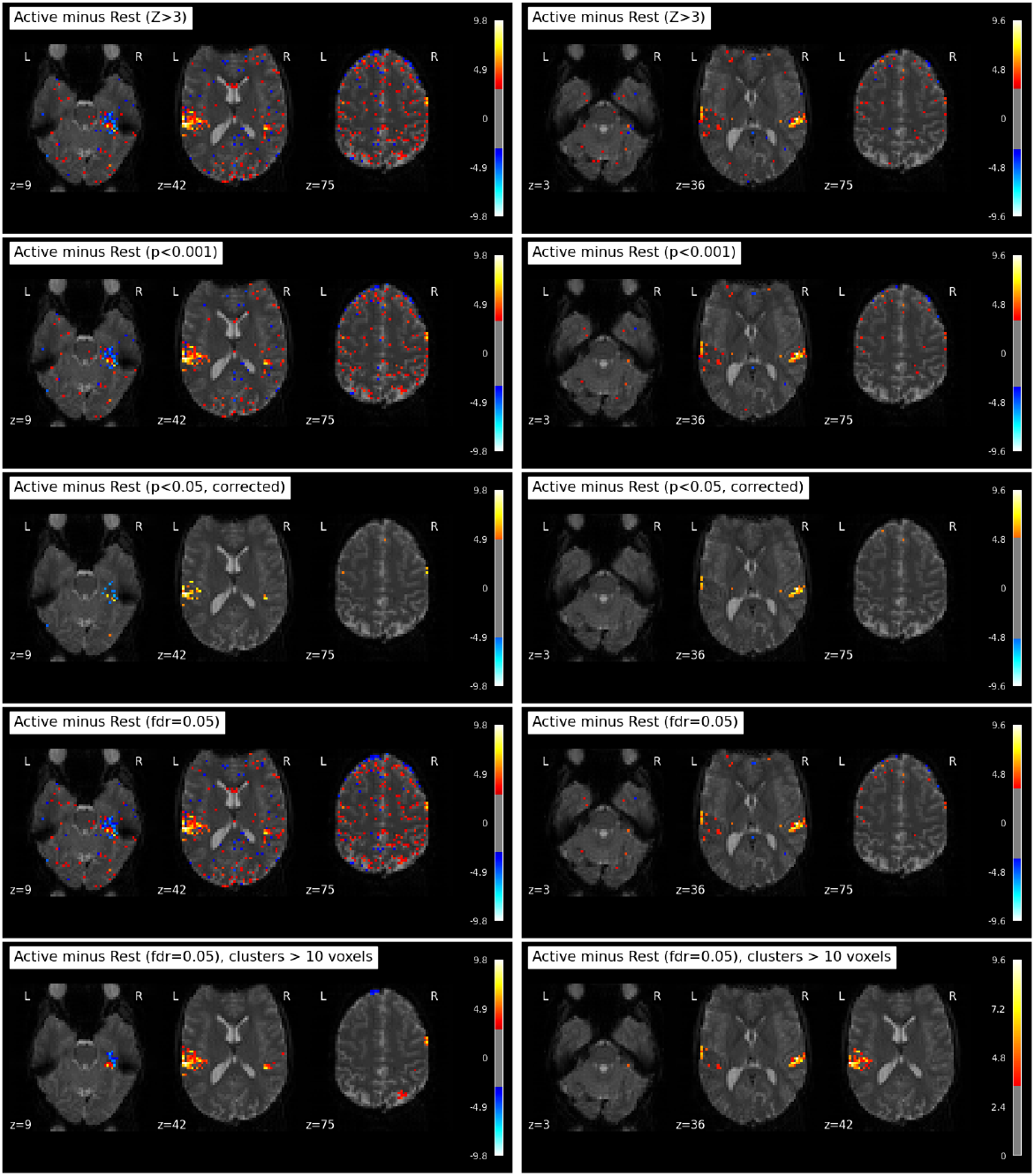
Left column: Standard GLM. Right column: GLM on reduced input.

- (*Z >* 3) test.
- (*p <* 0.001) test.
- Bonferroni corrected (*p <* 0.05) test.
- FDR = 0.05 corrected test.
- FDR = 0.05 corrected with cluster filtering test.

In the GLM representation shown in Fig. **1** (Left)(4.3), there is prominent red-highlighted activity in the auditory cortex, aligning with expectations. However, the model also detects signals in the parietal and frontal lobes Purves et al. (2001), located near the top of the head. These signals are likely extraneous, unrelated to auditory processing. Additionally, the GLM reveals various resting-state signals, depicted in blue. Notably, it sporadically identifies resting-state noise in both the Parietal and Frontal lobes, again seemingly unrelated to auditory functions. Intriguingly, there are signals near the auditory cortex and along the corticospinal tract, as shown in the leftmost column of Fig. **1** (4.3). One of the key challenges here is differentiating resting-state activity from movements in the corticospinal fluid, as they can easily be mistaken for one another.

We shift our attention the SIR adjusted GLM in Fig. **1** (right-column)(4.3) and note that across each statistical test the false activation in the parietal and frontal lobes is much smaller and is not detected at all in the FDR = 0.05 with cluster filtering test (Fig. **1** (4.3) bottom right-corner only picks up additional active-state signals near the left auditory cortex). This model also appears to favor detecting active state signals near the auditory cortex across all statistical tests. This suggests that the SIR model is detecting signals near relevant regions of interest and may even have the potential to detect appropriate BOLD responses in areas of the brain that tend to have more noise due to corticospinal fluid (i.e. the pyramidal tracts) since we don’t appear to detect signals associated with the corticospinal fluid.

Upon comparing the cluster formations in both models, we observe a distinct difference. The standard GLM model identifies nine clusters, as detailed in Table 1, with the largest cluster spanning a volume of 4050 mm^3^. In contrast, the SIR-adjusted GLM model, shown in Table 2, reveals only four clusters, with the largest being notably smaller at 2538 mm^3^. This observation aligns with the patterns typically highlighted with the auditory cortex Purves et al. (2001). The SIR model demonstrates a focused approach, predominantly recognizing clusters in proximity to the auditory cortex. Meanwhile, the standard GLM model exhibits a broader detection range, identifying signals in various brain regions, not exclusively linked to auditory processing.

**Table 1:**
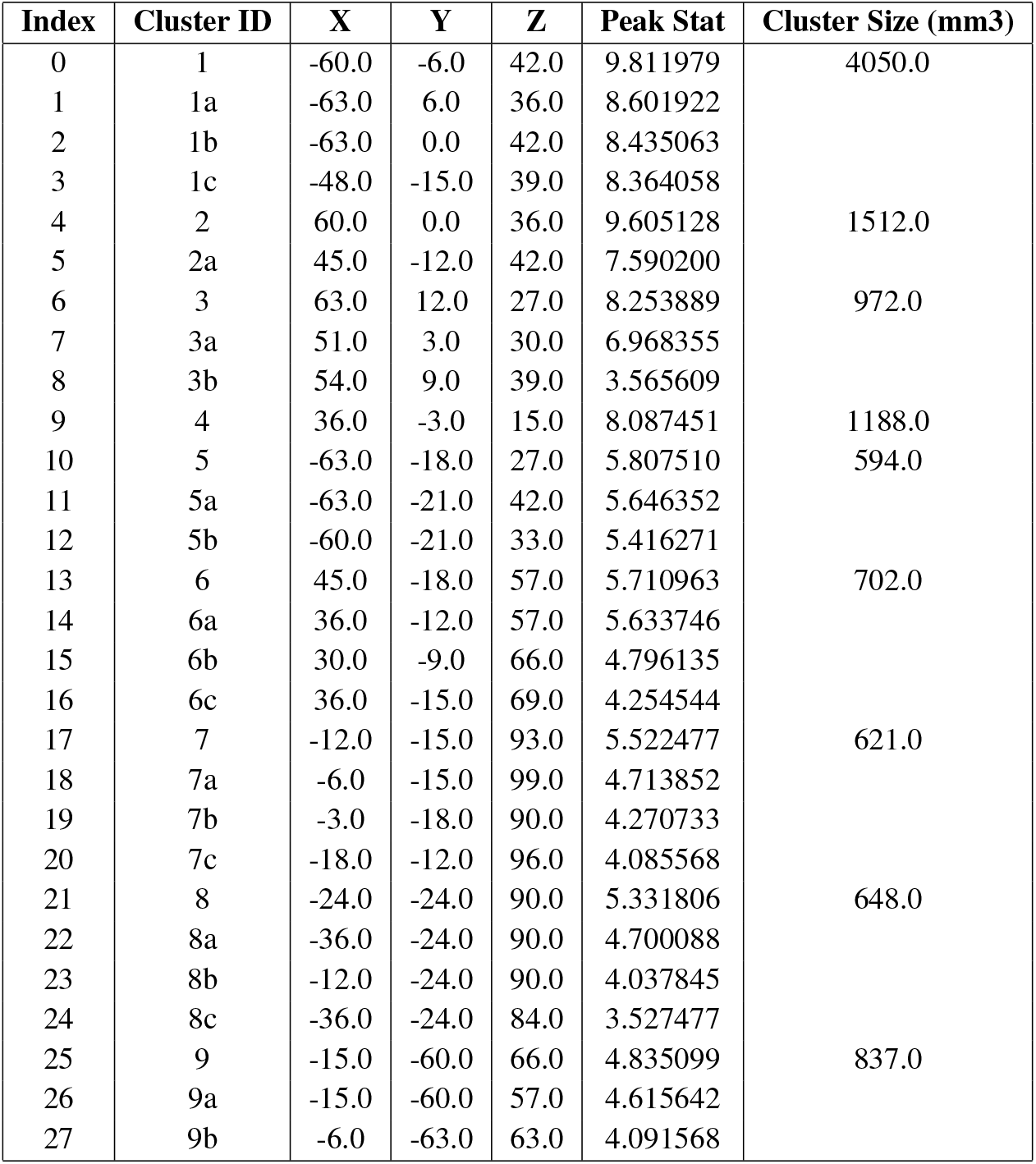
Detected Clusters (GLM Only)

**Table 2:**
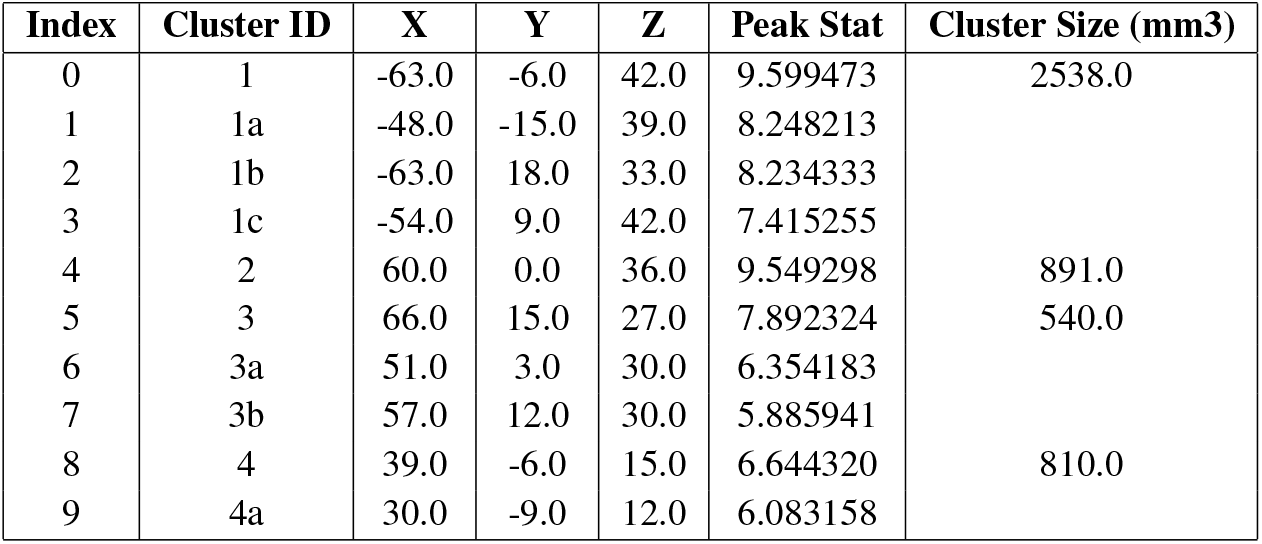
Detected Clusters (GLM + SIR)

## 4 Discussion

### 4.1 Summary of Findings

This research has shown the efficacy of Sliced Inverse Regression (SIR) in improving the accuracy and reliability of fMRI data analysis. By applying SIR to BOLD regressors, our study has demonstrated enhanced signal detection, reduced noise, and minimized artifacts compared to using General Linear Models (GLM) alone. These findings align with the current understanding of brain responses in active states, particularly in auditory-associated regions, and validate the anatomical relevance of our results.

### 4.2 Significance of SIR in fMRI Analysis

The introduction of SIR in fMRI analysis has the potential for significant advancement in neuroimaging methodologies. The dimensionality reduction achieved through SIR simplifies the interpretation of fMRI data by efficiently isolating relevant regressors. This simplification is critical in a field where the complexity of data often poses challenges in distinguishing signal from noise. The application of SIR, as demonstrated in our study, provides a clearer, more accurate representation of brain activity, enhancing the fidelity of fMRI as a tool for understanding neural mechanisms.

### 4.3 Implications for Future Research

The findings of this study have broad implications for the future of neuroimaging research. The improved accuracy in identifying brain regions associated with specific tasks or stimuli can refine our understanding of brain functionality. Furthermore, the open-access analysis of SIR methods provided in this study paves the way for wider adoption and further exploration of this technique in diverse fMRI applications. Researchers can now explore new avenues in neuroscience with a tool that offers greater precision and reliability. Traditionally, the reliance on numerous regressors in GLM has been a necessary yet problematic aspect of fMRI analysis, often leading to increased noise and complex interpretations.

Our results suggests that the integration of SIR into the analytical pipeline can address these issues. The comparison between GLM applied to original and SIR-adjusted data illustrates the superiority of the latter in terms of noise reduction and signal clarity. This advancement represents a notable shift from conventional methods, offering a more streamlined and effective approach to fMRI data analysis. Moreso, we don’t need to use overly sophisticated SIR methods as used in previous studies in order to improve the GLM analysis which has the potential to simplify these studies.

Recent advances in geometric deep learning and probabilistic generative modelling offer a rich but as–yet-under-explored toolkit for functional MRI. Geometry-aware autoencoders suggested in Lizarraga et al. (2022) and differentiable VQ encoders for tractography Lizarraga et al. (2024) show that respecting Wasserstein and quantization geometry yields compact yet faithful representations of highly heterogeneous anatomical data. Transferring these ideas to the spatio-temporal domain of BOLD could enable subject-specific latent codes that remain comparable across scanners and cohorts.

Registration methods that treat streamlines as points on a Riemannian manifold—e.g. deformation transfers transport Lizarraga et al. (2021) or diffeomorphic functional alignment of SrvfNet Nunez et al. (2021) suggest principled ways to align voxel-wise responses while preserving intrinsic temporal dynamics. From a probabilistic standpoint, PAC-Bayesian diffusion models for inverse problems Jiang et al. (2024) and MCMC-driven refinements Zhu et al. (2024) provide theoretically grounded regularizers that could be embedded directly into SIR pipelines to improve both detection power and uncertainty quantification.

Complementary insights arise from approaches that explicitly monitor *primitive interactions* during deep-network training Ren et al. (2025), offering a route to diagnose when learned representations deviate from plausible neuro-physiology. Finally, planning-as-inference frameworks such as the Latent Plan Transformer Kong et al. (2024) and the Latent Adaptive Planner Noh et al. (2025) demonstrate how hierarchical latent spaces can capture long-range temporal structure; adapting these transformers to multi-trial fMRI may bridge the gap between BOLD fluctuations and higher level cognitive sequences.

Taken together, these emerging techniques outline a potentially impactful research agenda for more expressive, interpretable, and biologically grounded models of BOLD activation.

## References

Alexandre Abraham, Fabian Pedregosa, Michael Eickenberg, Philippe Gervais, Andreas Mueller, Jean Kossaifi, Alexandre Gramfort, Bertrand Thirion, and Gaël Varoquaux. Machine learning for neuroimaging with scikit-learn. Front Neuroinform, 8:14, February 2014.

R. Coudret, S. Girard, and J. Saracco. A new sliced inverse regression method for multivariate response. Computational Statistics & Data Analysis, 77:285–299, 2014. ISSN 0167-9473. doi: 10.1016/j.csda.2014.03.006. URL https://www.sciencedirect.com/science/article/pii/S0167947314000838.

Maria Luisa Gorno-Tempini, Chloe Hutton, Oliver Josephs, Ralf Deichmann, Cathy Price, and Robert Turner. Echo time dependence of bold contrast and susceptibility artifacts. NeuroImage, 15(1):136–142, 2002. ISSN 1053-8119. doi: 10.1006/nimg.2001.0967. URL https://www.sciencedirect.com/science/article/pii/S105381190190967X.

Saad Jbabdi. Chapter 25 -imaging structure and function. In Heidi Johansen-Berg and Timothy E.J. Behrens (eds.), Diffusion MRI (Second Edition), pp. 585–605. Academic Press, San Diego, second edition edition, 2014. ISBN 978-0-12-396460-1. doi: 10.1016/B978-0-12-396460-1.00025-1. URL https://www.sciencedirect.com/science/article/pii/B9780123964601000251.

Eric Hanchen Jiang, Yasi Zhang, Zhi Zhang, Yixin Wan, Andrew Lizarraga, Shufan Li, and Ying Nian Wu. Unlocking the potential of text-to-image diffusion with pac-bayesian theory, 2024. URL https://arxiv.org/abs/2411.17472.

Deqian Kong, Dehong Xu, Minglu Zhao, Bo Pang, Jianwen Xie, Andrew Lizarraga, Yuhao Huang, Sirui Xie, and Ying Nian Wu. Latent plan transformer for trajectory abstraction: Planning as latent space inference. In A. Globerson, L. Mackey, D. Belgrave, A. Fan, U. Paquet, J. Tomczak, and C. Zhang (eds.), Advances in Neural Information Processing Systems, volume 37, pp. 123379–123401. Curran Associates, Inc., 2024. URL https://proceedings.neurips.cc/paper_files/paper/2024/file/df22a19686a558e74f038e6277a51f68-Paper-Conference.pdf.

Ker-Chau Li. Sliced inverse regression for dimension reduction. Journal of the American Statistical Association, 86(414):316–327, 1991. ISSN 01621459. URL http://www.jstor.org/stable/2290563.

Andrew Lizarraga, David Lee, Antoni Kubicki, Ashish Sahib, Elvis Nunez, Katherine Narr, and Shantanu H. Joshi. Alignment of tractography streamlines using deformation transfer via parallel transport. In Suheyla Cetin-Karayumak, Daan Christiaens, Matteo Figini, Pamela Guevara, Noemi Gyori, Vishwesh Nath, and Tomasz Pieciak (eds.), Computational Diffusion MRI, pp. 96–105, Cham, 2021. Springer International Publishing. ISBN 978-3-030-87615-9.

Andrew Lizarraga, Katherine L. Narr, Kirsten A. Donals, and Shantanu H. Joshi. Streamnet: A wae for white matter streamline analysis. In Erik Bekkers, Jelmer M. Wolterink, and Angelica Aviles-Rivero (eds.), Proceedings of the First International Workshop on Geometric Deep Learning in Medical Image Analysis, volume 194 of Proceedings of Machine Learning Research, pp. 172–182. PMLR, 18 Nov 2022. URL https://proceedings.mlr.press/v194/lizarraga22a.html.

Andrew Lizarraga, Brandon Taraku, Edouardo Honig, Ying Nian Wu, and Shantanu H. Joshi. Differentiable vq-vae’s for robust white matter streamline encodings. In 2024 IEEE International Symposium on Biomedical Imaging (ISBI), pp. 1–5, 2024. doi: 10.1109/ISBI56570.2024.10635543.

Torben E. Lund, Kristoffer H. Madsen, Karam Sidaros, Wen-Lin Luo, and Thomas E. Nichols. Non-white noise in fmri: Does modelling have an impact? NeuroImage, 29(1):54–66, 2006. ISSN 1053-8119. doi: 10.1016/j.neuroimage.2005.07.005. URL https://www.sciencedirect.com/science/article/pii/S105381190500501X.

Martin M Monti. Statistical analysis of fMRI Time-Series: A critical review of the GLM approach. Front Hum Neurosci, 5:28, March 2011.

Donghun Noh, Deqian Kong, Minglu Zhao, Andrew Lizarraga, Jianwen Xie, Ying Nian Wu, and Dennis Hong. Latent adaptive planner for dynamic manipulation, 2025. URL https://arxiv.org/abs/2505.03077.

Elvis Nunez, Andrew Lizarraga, and Shantanu H. Joshi. Srvfnet: A generative network for unsupervised multiple diffeomorphic functional alignment. In Proceedings of the IEEE/CVF Conference on Computer Vision and Pattern Recognition (CVPR) Workshops, pp. 4481–4489, June 2021.

D Purves, GJ Augustine, and et al. D Fitzpatrick. Neuroscience; The Auditory Cortex. Sunderland (MA): Sinauer Associates; https://www.ncbi.nlm.nih.gov/books/NBK10900, 2001.

Mehrdad Razavi, Thomas J Grabowski, Walter P Vispoel, Patrick Monahan, Sonya Mehta, Brent Eaton, and Lizann Bolinger. Model assessment and model building in fmri. Human brain mapping, 20(4), 2003–12. ISSN 1065-9471.

Jie Ren, Xinhao Zheng, Jiyu Liu, Andrew Lizarraga, Ying Nian Wu, Liang Lin, and Quanshi Zhang. Monitoring primitive interactions during the training of dnns. Proceedings of the AAAI Conference on Artificial Intelligence, 39(19):20183–20191, Apr. 2025. doi: 10.1609/aaai.v39i19.34223. URL https://ojs.aaai.org/index.php/AAAI/article/view/34223.

Y.H. Tu, Z.N. Fu, A. Tan, G. Huang, L. Hu, Y.S. Hung, and Z.G. Zhang. A novel and effective fmri decoding approach based on sliced inverse regression and its application to pain prediction. Neurocomputing, 273:373–384, 2018. ISSN 0925-2312. doi: 10.1016/j.neucom.2017.07.045. URL https://www.sciencedirect.com/science/article/pii/S0925231217313474.

Yiheng Tu, Ao Tan, Zening Fu, Yeung Sam Hung, Li Hu, and Zhiguo Zhang. Supervised nonlinear dimension reduction of functional magnetic resonance imaging data using sliced inverse regression. In 2015 37th Annual International Conference of the IEEE Engineering in Medicine and Biology Society (EMBC), pp. 2641–2644, 2015. doi: 10.1109/EMBC.2015.7318934.

Yaxuan Zhu, Zehao Dou, Haoxin Zheng, Yasi Zhang, Ying Nian Wu, and Ruiqi Gao. Think twice before you act: Improving inverse problem solving with mcmc, 2024. URL https://arxiv.org/abs/2409.08551.

